# Neural Progenitors as a Novel Pathogenic Mechanism in Microcephaly

**DOI:** 10.1101/2025.08.12.669854

**Authors:** Rami Yair Tshuva, Jeyoon Bok, Mio Nonaka, Xufeng Xue, Bidisha Bhattacharya, Aditya Kshirsagar, Tsviya Olender, Miri Danan-Gotthold, Tamar Sapir, Jianping Fu, Orly Reiner

## Abstract

Despite their significance, the genetic and molecular bases of neurodevelopmental disorders remain poorly understood. In this study, using human brain organoids and mouse models, we show that loss of *NDE1*, a gene closely associated with microcephaly, disrupts progenitor identity, prolongs mitosis, and alters regional patterning in the forebrain. *NDE1* knockout leads to a caudal identity shift of neural progenitor cells in the organoids and mouse brains, coinciding with aberrant ERK signaling. Notably, downstream activation of the ERK pathway restored rostral *PAX6* expression in human brain organoids. Parallel analyses of *Nde1* knockout mice confirmed disrupted regional patterning of the forebrain. Together, our data establish *NDE1* as a critical regulator of early human brain regionalization and elucidate molecular mechanisms underlying the structural abnormalities observed in *NDE1*-associated microcephaly.

## Introduction

Neurodevelopmental disorders are among the most pressing challenges in modern medicine, yet their genetic and molecular bases remain poorly understood^1–4^. Recent stem cell-based models offer powerful experimental tools to dissect these disorders in human-relevant systems. Microcephaly is a particularly severe group of neurodevelopmental disorders marked by profound abnormalities in brain size and structure, including microlissencephaly, characterized by a small, smooth brain, and microhydranencephaly, in which most of the cerebral hemispheres are replaced by a liquid-filled, membranous structure^5–13^. One gene linked to microcephaly is *NDE1*, which encodes the NDE1 protein believed to act in concert with LIS1 to mediate cellular activities critical for brain development, including neuronal migration, mitotic spindle orientation, and RNA regulation^5–29^. Besides microcephaly, *NDE1* mutations have been associated with a range of other neurodevelopmental and psychiatric disorders, including schizophrenia, intellectual disabilities, and autism spectrum disorders^9–11,30–42^. Thus, understanding how *NDE1* mutations disrupt early brain development is critical to identifying the shared mechanisms that contribute to the wide range of complex phenotypes observed in neurodevelopmental disorders.

## Results

To explore the roles of *NDE1*, we first examined its expression in the first-trimester human developing brain^43^ (Fig. 1a-f). *NDE1* is expressed in cycling progenitors, and its expression peaks during the G2/M state, similar to Aurora-kinase (AURKA), a typical marker of this state. To examine the function of *NDE1*, we generated *NDE1* knockout (KO) human pluripotent stem cell (hPSC) lines (Fig. 1g), and the cell lines were differentiated into brain organoids using an unguided on-chip protocol^44^ (Methods; Fig. 1h-l). The *NDE1* KO on-chip organoids displayed a reduced growth rate and decreased folding compared to wild-type (WT) controls (Fig. 1h-j). These findings are consistent with our previous work demonstrating that mutations in LIS1, a gene associated with lissencephaly, lead to diminished cortical folding^17,44^. Time-lapse imaging with H2B-mcherry and Lyn-GFP hPSCs enabled the analysis of specific phases of mitosis during brain organoid development (Fig. 1k-l). Overall, *NDE1* KO organoids exhibited a protracted duration of mitosis, with significant effects on prophase, metaphase, and telophase (Fig. 1l). Our findings fit the notion that many microcephaly-associated genes regulate the cell cycle^45–47^.

**Fig. 1.**
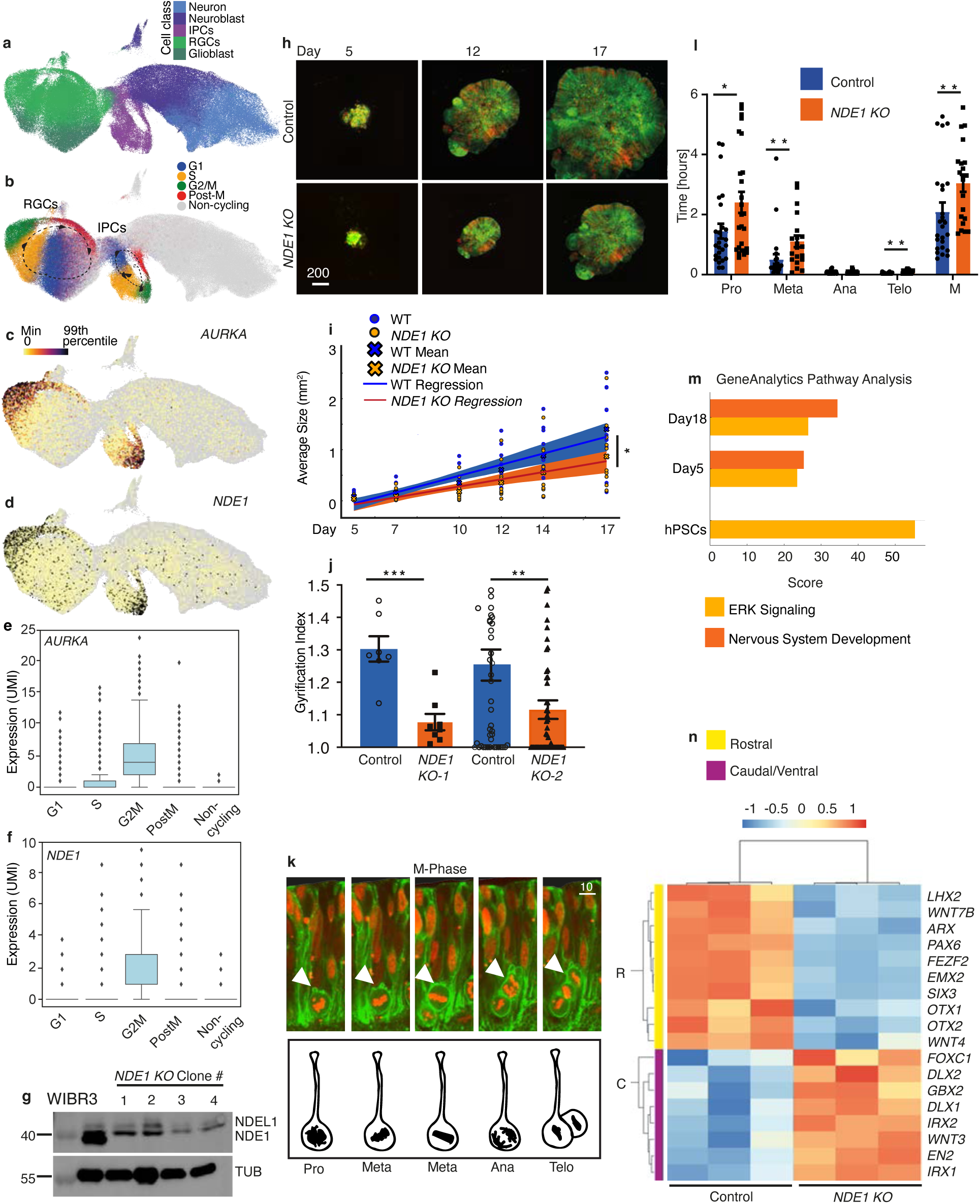
Characterization of *NDE1* KO human on-chip brain organoids reveals disrupted progenitor identity and altered ERK signaling. (**a-d**) UMAP visualization of human cortical excitatory neuron lineage (EMX2-positive) (taken from Braun et al.^43^), colored by (**a**) major cell classes (RGCs, radial glia cells, IPCs, intermediate progenitor cells). (**b**) proliferation state (G1, S, G2M, post-M), non-cycling cells are colored in grey, (**c**) expression patterns of *AURKA* and (**d**) *NDE1,* note that both genes are highly expressed in cells at the G2/M state. (**e** and **f**) Box plot of *AURKA* (**e**) and *NDE1* (**f**) expression across proliferation states (only v3 chemistry samples, see Methods). (**g**) Representative western blot showing the absence of NDE1 four *NDE1* KO clones used in this study. Tubulin serves as a loading control. (**h**) Representative images of WT and *NDE1* KO on-chip organoids at indicated differentiation days (5, 12, 17), labeled with H2B-mCherry (red) and Lyn-GFP (green). Scale bar, 200 µm. (**i**) Organoid size quantification (n ≥8 organoids per group, two biological repeats; Growth curves were analyzed by linear regression of organoid diameter versus time, comparing slopes using ANCOVA. Mean values±SEM are presented, slopes were significantly different, P=0.0418*). (**j**) *NDE1* KO organoids exhibit fewer folds than the control, indicated by the gyrification index (mean ± SEM, WT n=7, *NDE1* KO-1 n=8, P=0.0006***, WT n=47, *NDE1* KO-2 n=55, P=0.042**, Mann–Whitney test). Each n represents a single organoid (**k**) Fluorescent time-lapse images of a dividing cell (indicated by the white arrowheads). The illustration below shows prophase (Pro), metaphase (Met), anaphase (Ana), and telophase (Tel). Scale bar, 10 µm. (**l**) Quantification of the mitosis (M) duration according to the indicated stages (mean ± SEM, WT n=20 cells, *NDE1* KO n=25 cells, Mann-Whitney test, prophase P=0.0276*, metaphase P=0.0013**, telophase P=0.0074**). (**m**) Histogram comparing GeneAnalytics scores for "ERK Signaling" Super Path and "Nervous System Development" pathways across developmental stages (hPSCs, Day 5, and Day 18). (**n**) Heatmap illustrating changes in regional marker gene expression across the rostrocaudal and dorsoventral axes in WT and *NDE1* KO organoids (color scale indicates normalized expression levels, n = 3 organoids per sample).

We conducted bulk RNA-seq for the *NDE1* KO and WT hPSCs and the on-chip brain organoids derived from these cells on days 5 and 18 (Supplementary Data S1). Gene Analytics^48^ analysis of differentially expressed genes (DEGs) revealed robust enrichment for "ERK Signaling" in the *NDE1* KO cells across developmental stages, peaking at the pluripotent stage and remaining elevated in brain organoids (Fig. 1m). Enrichment for "Nervous System Development" emerged at Day 5 and became more notable on Day 18, reflecting progressive neural differentiation and maturation (Fig. 1m). Further examination of DEGs revealed that in *NDE1* KO organoids, rostral genes, including *EMX2*, *LHX2*, and *PAX6*, were down-regulated, whereas caudal and ventral genes, including *EN2* and *DLX1*, were up-regulated, suggesting a shift in the regional identity of neural progenitors in the organoids (Fig. 1n). These results were corroborated using real-time PCR (Supplementary Fig. S1a, Supplementary Table S1). Consistently, mouse models with reduced ERK signaling exhibit microcephaly^49,50^, and furthermore, ERK signaling regulates PAX6 expression in the developing CNS^49^.

Considering that the on-chip organoids are not patterned to a specific brain region identity, we sought additional strategies to examine the function of *NDE1* in brain patterning. To this end, we generated hPSC-derived organoids with either a cortical or cerebellar identity^51,52^ (Methods; Supplementary Fig. S1b-h). *NDE1* KO cortical and cerebellar organoids both exhibited smaller sizes than WT controls (Supplementary Fig. S1b). However, the size disparity resulting from *NDE1* KO was more pronounced for cortical organoids. The smaller organoid size was not due to increased apoptosis, premature differentiation, or cell cycle exit (Supplementary Fig. S1c-h). Instead, phospho-histone immunostaining pointed toward differences in mitosis, consistent with previous studies^53^ (Supplementary Fig. S1c-h).

We then conducted scRNA-seq for both *NDE1* KO and WT cortical organoids. A significant reduction in cells with a forebrain identity was noted in *NDE1* KO cortical organoids compared to WT controls (Fig. 2a, Supplementary Fig. S2a-g). When the regional identity of each cell was quantified, a significant decrease in the ratio of cells with forebrain, telencephalon, and diencephalon identities and an increase in the ratio of cells exhibiting the identities of midbrain, hindbrain, medulla, pons, and cerebellum were noted (Fig. 2b, Supplementary Fig. S2a-g). These regional fate identity changes were corroborated using real-time PCR, with *NDE1* KO cortical organoids exhibiting reduced expression of rostral genes, including *TBR1*, *EMX2*, and *FOXG1,* but upregulating caudal genes *EN1*, *EN2*, *PAX2*, and *IRX2* (Supplementary Fig. S2h). These convergent gene expression data support that *NDE1* loss disrupts early rostrocaudal brain patterning, diverting progenitors away from rostral forebrain fates and toward more caudal mid/hindbrain and cerebellar identities.

**Fig. 2.**
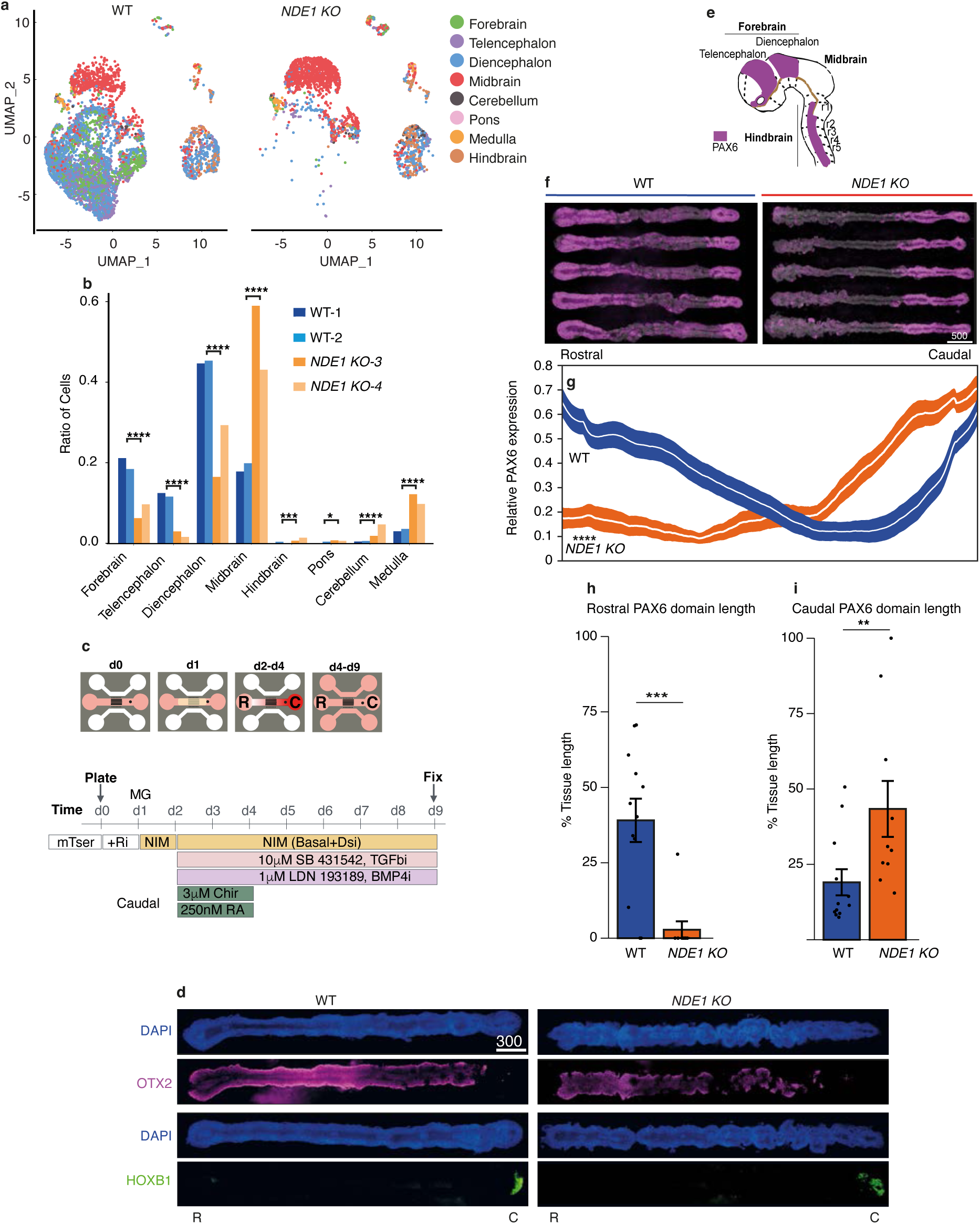
Single-cell RNA analysis identifies altered progenitor identity and signaling. **(a)** UMAP of scRNA-Seq data of cortical organoids (Day 30). The regions are color-coded as indicated based on the human embryonic brain atlas^43^. (**b**) Barplot showing the ratio of WT and *NDE1* KO organoid cells mapped to each region (significance was determined by two-sided Fisher’s exact test, *P<0.05, ***P<0.001, ****P<0.0001) (**c**) Schematic representation of rostrocaudal patterning strategy used for neural morpho chip (NMC) differentiation, including experimental timeline. (**d**) Representative immunofluorescence images comparing regional marker protein expression (OTX2, HOXB1) combined with DAPI in WT and *NDE1* KO NMC. Rostral, R, and Caudal, C, sides are indicated (**scale bar: 300 µm**). (**e**) Schematic depicting PAX6 expression pattern in the rostral (forebrain, magenta) and caudal (hindbrain, magenta) domains. (**f**) Representative immunofluorescence images showing altered spatial distribution of PAX6-positive cells (magenta) in WT and *NDE1* KO NMCs (scale bar, 500 µm). (**g**) Quantification of the distribution of PAX6-positive cells along the rostrocaudal axis (mean ± SEM, n = 52 WT organoids and n = 36 KO organoids, functional ANOVA, permutation test). (**h**, **i**) Quantification of rostral (**h**) and caudal (**i**) PAX6 domain lengths in WT and *NDE1* KO NMCs, expressed as percentage of total organoid length (mean ± SEM, n = 13 WT NMCs and n = 10 KO NMCs; **j**, P =0.00032***, **k**, P=0.02894**, *Student’s t-test*).

Current brain organoids usually do not recapitulate axial neural patterning within each individual structure. To address this limitation, we leveraged a recently developed NeuroMorphoChip (NMC)^54^ system that applies microfluidic gradients for controlled formation of patterned neural organoids mimicking the entire CNS. By carefully adjusting microfluidic morphogen gradients, patterned forebrain-, midbrain-, and hindbrain-like regions could be induced in NMC organoids along their rostrocaudal axes (Methods; Fig. 2c,d). To investigate the role of *NDE1* in rostrocaudal neural patterning, immunostaining for PAX6 was conducted on both WT and *NDE1* KO NMC organoids. PAX6 is expressed in neuronal progenitors of the forebrain (covering telencephalon and diencephalon) and the hindbrain but is absent in the midbrain (Fig. 2e)^55^. In WT NMC tissues, PAX6 exhibited high expression in the rostral region corresponding to the forebrain, minimal expression in the mid-section associated with the midbrain, and moderate expression in the caudal, hindbrain-like area (Fig. 2f-i). In contrast, *NDE1* KO NMC organoids showed pronounced reduction in rostral PAX6 expression, accompanied by a notable expansion of caudal PAX6+ domains (Fig. 2f-i). This altered spatial patterning of PAX6 expression underscores a critical role for *NDE1* in the proper regionalization of neural progenitors along the rostrocaudal axis.

scRNA-seq of the rostral halves of WT and *NDE1* KO NMC organoids further revealed a distinct cell cluster missing due to *NDE1* KO (Fig. 3a-c). This cluster is enriched for genes associated with signal transduction, cell cycle, mitosis, and ERK signaling (Fig. 3d, Supplementary Fig. S3a-l). When projected onto the data of radial glia from PCW 5 and 5.5 human brain atlas, the major progenitor population at this developmental stage, cells in the rostral halves of both WT and *NDE1* KO NMC tissues overlapped mainly with the forebrain radial glial cluster (Fig. 3e-g). Furthermore, the missing cell population in *NDE1* KO NMC organoids was also mainly projected to the forebrain region (Fig. 3h,i). Together, these data support that the rostral regions of *NDE1* KO NMC organoids were depleted of a subset of forebrain cells enriched with ERK signaling-related genes.

**Fig. 3.**
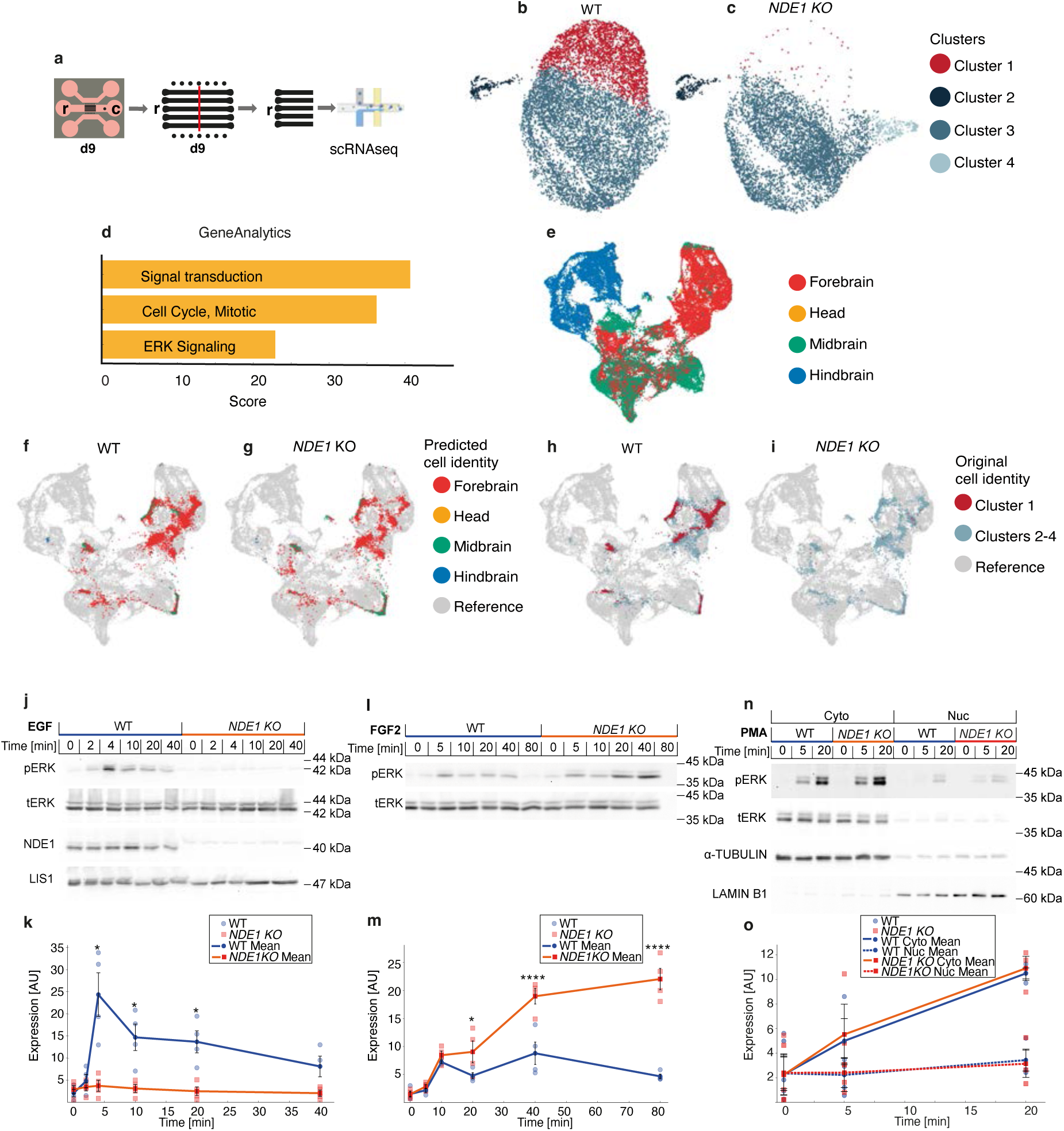
scRNA-seq data and altered ERK signaling dynamics and regional identity in *NDE1* KO cells and NMCs. (**a**) Schematic representation of sample preparation for single-cell RNA sequencing. Tissues were cut in half, and the rostral halves were further processed. (**b**) UMAP revealed a missing cluster (cluster 1) in scRNA seq of the rostral halves of *NDE1* KO (**c**) compared to WT NMC’s (**b**). (**d**) GeneAnalytics analysis of cluster 1 **(e-i)** UMAP visualization of 5 and 5.5 PCW radial glia cells from the human embryonic brain atlas^43^, color-coded by anatomical region, (**f,g**) or the cluster identity **(h,i)** from **b** and **c**. ERK signaling dynamics (**j-o**). (**j**, **l**, **n**) Representative western blots showing phosphorylated ERK (pERK) levels after stimulation with (**j**) EGF, (**l**) FGF2, and (**n**) PMA in WT and *NDE1* KO neural stem cells (NSCs). Total ERK (tERK) and β-actin were used as loading controls. (**k,m,o**) Quantification of western blot results from (**j**, **l**, **n**), respectively (mean ± SEM, n = 4 independent experiments, **b**, P<0.05*, **d**, P<0.00001****, **f**, all comparisons non-significant, Two Way ANOVA, posthoc Šídák’s multiple comparisons test.

Our results point to dysregulated ERK signaling as a potential molecular mechanism underlying pathological brain patterning. To dissect ERK signaling in the context of *NDE1* deficiency, we first chose neural stem cells (NSCs) as a cellular model to investigate. Both *NDE1* KO and WT hPSCs were differentiated into NSCs, before these cells were stimulated with distinct ERK pathway activators, including EGF, FGF2, and phorbol 12-myristate 13-acetate (PMA) (Methods; Fig. 3j-o). Western blot analyses revealed that *NDE1* KO NSCs failed to mount a robust ERK phosphorylation response following EGF stimulation, in contrast to WT controls (Fig. 3j,k). Upon FGF2 treatment, ERK activation increased continuously in *NDE1* KO NSCs, as opposed to an attenuated activity in WT cells (Fig. 3l,m). Stimulation with PMA, a direct protein kinase C (PKC) activator that bypasses receptor-level inputs, restored ERK phosphorylation and nuclear translocation in *NDE1* KO NSCs to levels comparable to those observed for WT cells (Fig. 3n,o). Together, these data support that *NDE1* is required for upstream ERK pathway activation in response to specific morphogen signals. Nonetheless, *NDE1* KO might not impair the core signaling machinery downstream of MEK, including ERK activation and nuclear entry.

To corroborate the *in vitro* findings, we next generated *Nde1* KO mice and examined brain patterning (Methods; Fig. 4a-l), which was not reported in prior studies of *Nde1* KO mice^21^. Compared to WT controls, mutant mouse embryos exhibited a substantial decrease in PAX6 immunoreactivity in both the telencephalon and diencephalon (Fig. 4a-h,k). OTX2, typically upregulated in the forebrain and midbrain, was expressed in both WT and *Nde1* KO mouse brains (Fig. 4i-j,k). However, in WT mouse brains, OTX2 expression peaked in the diencephalon, while in *Nde1* KO ones it was maximal in the diencephalon (Fig. 4i,j,l). Thus, there is a notable caudal shift of the OTX2 expression peak to the midbrain due to *Nde1* KO.

**Fig. 4.**
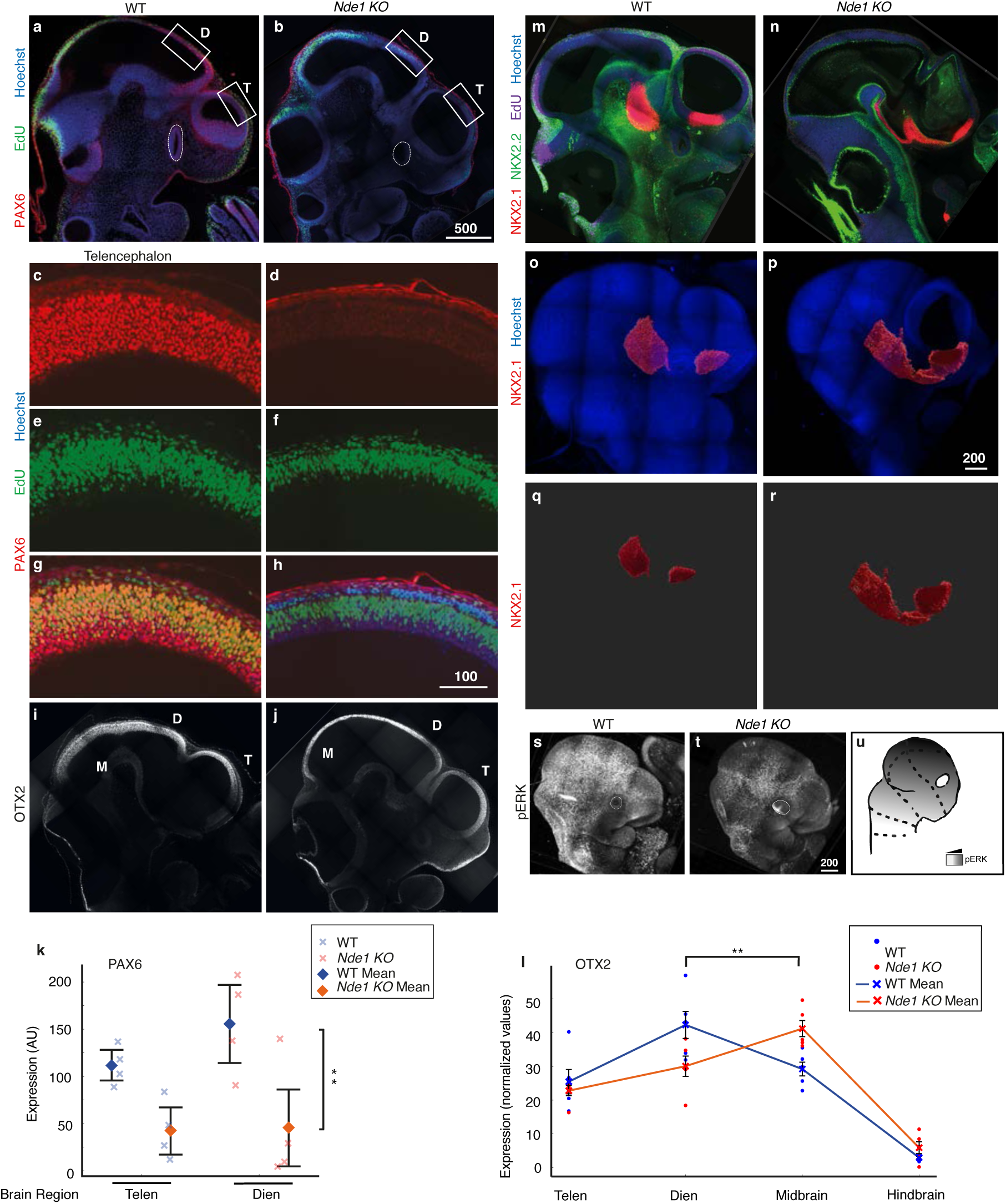
Altered brain regionalization and signaling in *Nde1* KO mouse embryos. (**a**, **b**) Immunofluorescence images of WT (**a**) and *Nde1* KO (**b**) Sagital optical Z slice from whole mount E10.5 mouse embryos stained for PAX6 (red), EdU (green), and Hoechst (blue); boxes indicate telencephalon (T) and diencephalon (D), the telencephalon regions are shown in higher magnification below **(scale bar: 500 µm**). (**c-h**) High magnification of boxed T regions in (**a**, **b**) showing individual channels: PAX6 (red, **c, d**), EdU (green, **e, f**), and merged (**g, h**). Scale bar**: 100 µm.** (**i**, **j**) Immunofluorescence of OTX2 expression highlighting telencephalon (T), diencephalon (D), and midbrain (M) in WT (**i**) and KO (**j**) embryos (**scale bar: 200 µm**). (**k**, **l**) Quantification of PAX6 (**k**) and OTX2 (**l**) normalized expression intensity in WT and *Nde1* KO embryos. PAX6 data (mean ± SEM; n = 4 embryos per genotype) was analyzed using two-way ANOVA, genotype-based difference is significant P=0.0019**. OTX2 peak distribution (normalized values mean±SEM; WT, n=5, *Nde1* KO n=6 embryos) was analyzed by Fisher’s exact test P=0.0022**. (**m-r**) Immunofluorescence images of NKX2.1 (red), NKX2.2 (green), EdU (magenta), and Hoechst (blue) in WT (**m**) and KO (**n**) embryos illustrating ventral patterning changes (scale bar: 200 µm). (**o-r**) *Nde1-/-* NKX2.1 staining (red) shown alone as a volume using IMARIS and merged with Hoechst (blue), highlighting altered ventral domains in WT (**o, q**) and KO (**p, r**) embryos. (**s-u**) Immunostaining for phosphorylated ERK (pERK) illustrating reduced ERK activation in *Nde1* KO (**t**) compared to WT (**s**) embryos; (**u**) shows a scheme of pERK expression pattern in the developing mouse embryonic brain.

Our on-chip brain organoids data revealed dysregulated ventral gene expression. We thus also examined two ventral markers in *Nde1* KO mouse brains: NKX2.2, which is primarily expressed in ventral forebrain, midbrain, and hindbrain, and NKX2.1, expressed in ventral forebrain, medial ganglionic eminence (MGE), and preoptic area (Fig. 4m-r)^56–58^. A striking difference was observed in NKX2.1 expression: while WT mouse brains displayed two distinct NKX2.1 expression domains, these areas appeared intermingled in mutant embryos (Fig. 4m-r).

Notably, in mouse brains, several regional identity markers - including FOXG1, PAX2, and PAX3 at the protein level, as well as *En1*, *Dlk1*, *Lmx1b*, and *Pax2* at the RNA level - remained largely unaffected by *Nde1* KO (Supplementary Fig. S3). These data suggest more subtle and specific patterning disruptions in mouse brains, as compared to the broader regional identity defects observed in the human models. Importantly, there was a decreased ERK activity, revealed by immunostaining for phosphorylated ERK, in *Nde1* KO mouse brains compared to WT controls (Fig. 4s-u).

We next examined whether ectopic activation of the ERK pathway could restore regional brain patterning in *NDE1* KO brain organoids. Given the importance of WNT inhibition in establishing forebrain identity^59^, we included the WNT inhibitor XAV939 (XAV). XAV treatment alone of *NDE1* KO NMC organoids increased rostral expression of PAX6 similarly to PMA, and combining XAV with PMA resulted in an additive enhancement of PAX6 expression (Fig. 5a). Our experimental approach, detailed in Fig. 5b, therefore incorporated rostral addition of XAV. In *NDE1* KO NMC tissues, rostral *PAX6* RNA expression was significantly reduced (Fig. 5c). However, PMA treatment restored rostral *PAX6* RNA expression to control levels. All rescue or control groups (*NDE1* KO + PMA, WT, WT + PMA, all with XAV) displayed significantly greater rostral expression of PAX6, as compared to untreated *NDE1* KO NMC organoids. Spatial quantification further confirmed a pronounced increase in PAX6+ cells in PMA-treated *NDE1* KO NMC organoids (Fig. 5e). Together, these data show that downstream activation of the ERK pathway is sufficient to partially rescue impaired rostral PAX6 expression in *NDE1*-deficient brain organoids. The roles of NDE1 and LIS1 in the FGF-ERK pathway have been established in a previous limb development study, which supports the positioning of NDE1 and LIS1 both upstream and downstream of FGF receptor signaling^60^.

**Fig. 5.**
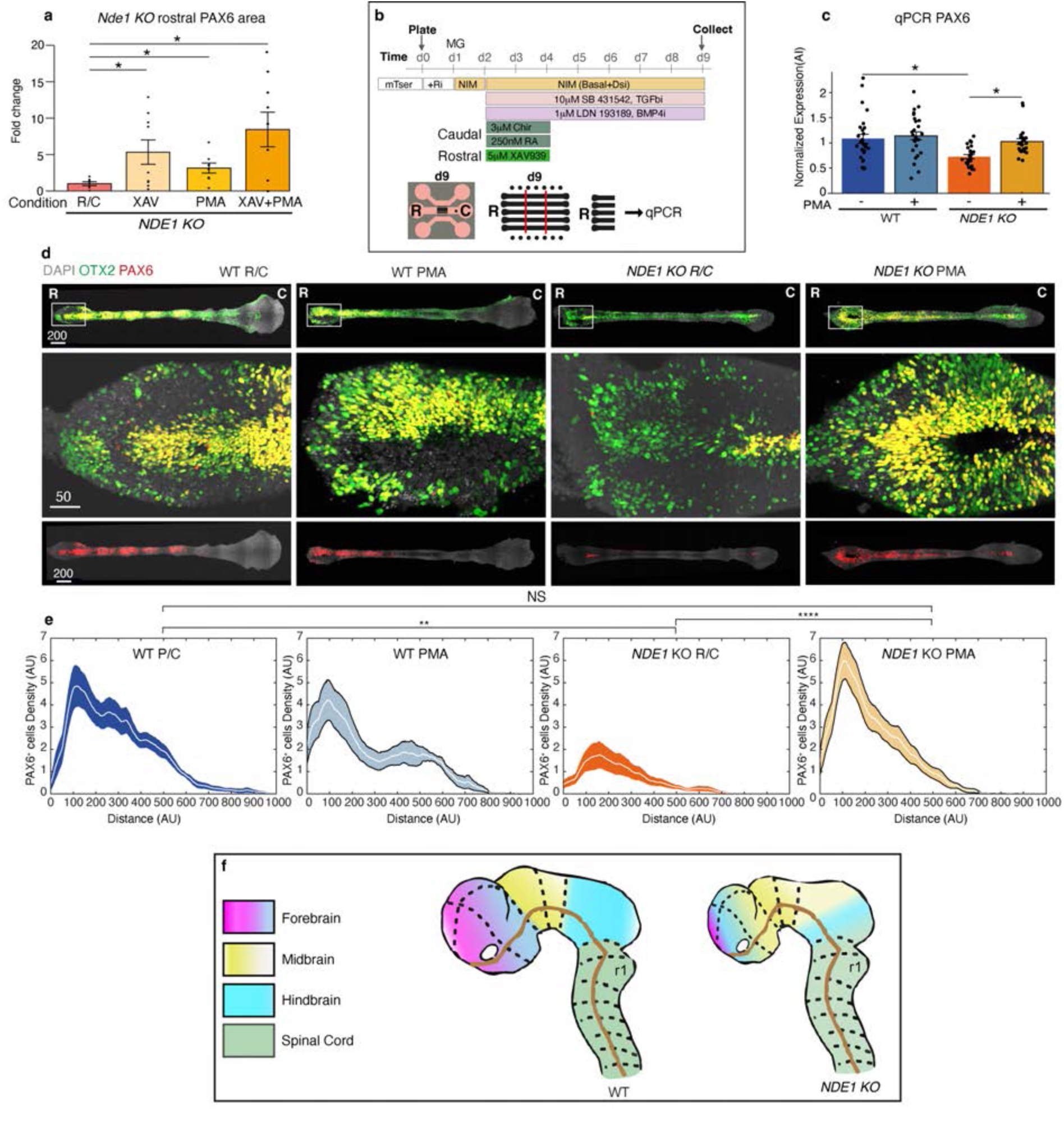
ERK pathway activation restores rostral PAX6 expression in *NDE1* KO organoids. (**a**) Expansion of PAX6 rostral domain in NMC following addition of either the WNT inhibitor (XAV939, XAV), PMA, or both, to the rostral media reservoir during patterning. (**b**) Experimental timeline and schematic for PMA and WNT inhibitor (XAV939, XAV) treatments in NMC organoids. (**c**) qPCR quantification of normalized PAX6 rostral expression across wild-type (WT) and *NDE1* KO conditions, with and without PMA treatment. Each point represents a technical replicate; bars show mean ± SEM of biological replicates. Biological samples WT n=5, WT+PMA n=8, *NDE1* KO n=5, *NDE1* KO+ PMA n=4. A linear mixed-effects model was applied with Genotype and Treatment as fixed effects and Biological Replicate as a random intercept. Significant differences were observed between WT and *NDE1* KO (p < 0.05), and between *NDE1* KO and *NDE1* KO+PMA (p < 0.05), indicating that PMA partially rescues PAX6 expression in NDE1 organoids. No significant interaction was found between Genotype and Treatment. (**d**) Representative immunofluorescence images showing expression of PAX6 (red), OTX2 (green), and DAPI (gray) in *NDE1* KO organoids with and without PMA treatment; magnified regions indicated by boxes (scale bars: 200µm for the top and bottom images and 50µm for the middle panel). (**e**) Quantification of the spatial distribution of PAX6-positive cells along the rostrocaudal axis in WT control (dark blue), WT+PMA (light blue), NDE1 KO (orange), and NDE1 KO+PMA (light orange) organoids. Shaded areas represent SEM; (WT n=9, WT + PMA n=7, *NDE1* KO n=9, *NDE1* KO + PMA n=10 patterned organoids. fANOVA, permutation test was used for statistical test, significance P=0****, P=0.002**, or NS, non-significant). (**f**) Schematic summarizing the regional identity shifts observed with *NDE1* KO in comparison to WT organoids (colors represent regions: pink = rostral forebrain; yellow = midbrain; blue = hindbrain; green = spinal cord).

In summary, this study unravels a previously unrecognized role of *NDE1* in regional patterning of neural progenitors in the brain. Using human brain organoids and mouse models, we show that loss of NDE1 impacts both brain size and regional identity (Fig. 5f). This patterning deficit was accompanied by dysregulated ERK signaling, which could be restored by downstream activation, rescuing PAX6 expression in *NDE1* KO brain organoids. Our data thus reveal a new mechanistic link between *NDE1*-mediated mitotic control and regional brain identity, mediated in part by ERK signaling associated with cytokines such as FGF and EGF. The rescue of rostral identity through ectopic activation of ERK signaling was also enhanced by WNT inhibition, suggesting cooperative interactions between these pathways in brain regional specification and demonstrating that dysregulation of brain regional patterning due to *NDE1* mutations might be partially reversible. Our data thus expand the classical view of NDE1 from being a “mitotic scaffold” that safeguards spindle orientation and progenitor pool size to a multifaceted regulator that also gates ERK signaling and, consequently, mediates brain regional patterning. Our data position NDE1 in the emerging category of microtubule-associated proteins as organizers of signaling complexes that coordinate proliferation dynamics with patterning signals, ensuring that daughter cells emerge with the correct regional identity rather than merely the correct number. The implications of this study are beyond just microcephaly, as ERK signaling might prove to be a viable therapeutic target in other neurodevelopmental disorders.

RNA-Seq data deposited

- MARSeq of on-chip organoids; To review GEO accession GSE232506: Go to https://www.ncbi.nlm.nih.gov/geo/query/acc.cgi?acc=GSE232506 Enter token cdqpqawmpbcjjav into the box
- scRNA-Seq of forebrain organoids; To review GEO accession GSE229988: Go to https://www.ncbi.nlm.nih.gov/geo/query/acc.cgi?acc=GSE229988 Enter token cdmfksoohjufben into the box
- scRNA-Seq of NeuroMorphoChip (NMC); To review GEO accession GSE279902: Go to https://www.ncbi.nlm.nih.gov/geo/query/acc.cgi?acc=GSE279902 Enter token sdgpamoanpwzryh into the box

## Acknowledgments

Orly Reiner is an incumbent of the Berstein-Mason professorial chair of Neurochemistry and the Head of the M. Judith Ruth Institute for Preclinical Brain Research. Tamar Sapir is the Incumbent of the Leir Research Fellow Chair in Autism Spectrum Disorder. We thank the transgenic mouse facility and the caretaker, Ms. Tamir Moshe, at the Weizmann Institute. Activities in the Fu lab have been supported technically by the Michigan Medicine Microscopy Core for microscopy imaging, the Michigan Advanced Genomics Core for scRNA-seq service, and the Michigan Lurie Nanofabrication Facility for microfabrication.

## Funding

Israel Science Foundation ISF grant (545/21)

United States-Israel Binational Science Foundation (BSF; Grant No. 2023009)

NSF-BSF Emerging Frontiers in Research and Innovation (EFRI) (NSF-BSF; Grant No. 2024616)

Israel Ministry of Innovation, Science and Technology IL (0005900) Azrieli Institute for Brain and Neural Sciences

The Maurice and Vivienne Wohl Biology Endowment

The Gladys Monroy and Larry Marks Center for Brain Disorders The Advantage Trust

The Nella and Leon Benoziyo Center for Neurological Diseases

The David and Fela Shapell Family Center for Genetic Disorders Research The Abish-Frenkel RNA center

The Andrea L. and Lawrence A. Wolfe Family Center for Research on Neuroimmunology and Neuromodulation,

The Weizmann Center for Research on Neurodegeneration The Brenden-Mann Women’s Innovation Impact Fund

The Irving B. Harris Fund for New Directions in Brain Research

The Irving Bieber, M.D., and Toby Bieber, M.D. Memorial Research Fund

The Leff Family, Barbara & Roberto Kaminitz, Sergio & Sônia Lozinsky, Debbie Koren, Jack and Lenore Lowenthal, and the Dears Foundation

A research grant from the Estates of Ethel H. Smith, Gerald Alexander, Mr. and Mrs. George Zbeda, David A. Fishstrom, Norman Fidelman, Hermine Miller, Olga Klein Astrachan and Hermine Miller.

Ethel Lena Levy, the Selsky Memory Research Project

National Science Foundation of the United States (CBET 1901718, PFI 2213845, and EFMA 2422149)

National Institutes of Health of the United States (R21 NS127983, R01 GM143297, and R01 NS129850)

## Author contributions

Conceptualization: JF, TS, OR Methodology: RYT, JB, MN, XX, AK

Investigation: RYT, JB, MN, AK, BB, TS, MDG Visualization: RYT, JB, MN, BB, TS, MDG, TO Funding acquisition: JF, OR

Project administration: JF, TS, OR Supervision: XX, TO, MN, TS, JF, OR Writing – original draft: OR, TS, JF

Writing – review & editing: RYT, JB, MN, XX, BB, AK, TO, MDG, TS, JF, OR

## Data and materials availability

All data, code, and materials used in the analysis is available in some form to any researcher for purposes of reproducing or extending the analysis. Transfer of cell lines, mice, and plasmids will require materials transfer agreements (MTAs). RNA-seq accession numbers have been indicated.

## Supplementary Materials

Materials and Methods Figs. S1 to S4

Tables S1 to S2 References (*1-33*) Data S1

## Materials and Methods

### Ethics statement

Work with hESC (WIBR3, NIHhESC-10-0079) and genome editing was carried out with approval from the Weizmann Institute of Science IRB (Institutional Review Board). Work with animals was carried out with approval from the Weizmann Institute of Science IACUC (Institutional Animal Care and Use Committee). The use of experimental animals is in complete accordance with: The Animal Welfare Law (Experiments with animals); The Regulations of the Council for Experiments with Animals; The Weizmann Institute Regulations (SOP); The Guide for the Care and Use of Lab Animals, National Research Council, 8th edition; The Guidelines for the Care and Use of Mammals in Neuroscience and Behavioral Research.

### hPSC culture, genome editing of human and mouse lines

WIBR3 (NIHhESC-10-0079) hPSCs obtained from the Jaenisch lab^1^, and isogenic *NDE1-*KO were cultured on irradiated MEFs (mouse embryonic fibroblasts) in NHSM media as previously described^2,3^. Cells were routinely checked for Mycoplasma. Pluripotency was evaluated using FACS analysis of pluripotent markers (SSEA-4 PE and TRA-1-60 Alexa Fluor 647, >90% positive). Two different gRNA from coding exon 4 were used to generate the four different *NDE1-*KO lines. The gRNAs were introduced using pX330 by electroporation with the addition of a GFP expression plasmid ^4,5^. Three days after transfection, the cells were subjected to FACS and plated at a density of 2,000 cells per 10 cm plate on irradiated MEFs, allowing for the growth of single-cell-derived colonies. The mutation was confirmed by PCR, restriction enzyme digestion, Sanger DNA sequencing, and Western blot analysis. In the case of all four lines, no protein was detected (Fig. 1G). Line 1 (1G) was generated using gRNA (5’- GTTCCAGCTCCATGCGAAGG), Sanger sequencing revealed a deletion of 11bp (CCTTCGCATGG) and an addition of 1bp (A). The rest of the lines were generated by gRNA (5’-GGAACTCCGAGAATTCCAGG). Line 2 (42) had a deletion of 1bp (C), line 3 (13) had a deletion of 256bp, line 4 (51) had a deletion of 1bp (C).

The *Nde1* KO mice were generated by the Transgenic and Embryo Manipulation Lab at Weizmann, using the CRISPR/CAS9 system with two different guides on exon six: TCTACTCCAGTAGCTCACCG and CAGGCTCAGGGCAAGCCAAG. The mice were outbred to generate distinct lines. In most of the studies, line 51 was used; the mutation was an additional A, resulting in a premature stop codon and a short predicted protein of 82 amino acids. Line 2 (42) had a deletion of one C.

For size and folding evaluation, the actin cytoskeleton was stably labeled with Lifeact- or Lyn-GFP, and nuclei were labeled with H2B-mCherry PiggyBac plasmids^6,7^ together with the transposase, to generate stable control and isogenic KO lines, as previously described^3^.

### Western blot

Forty-eight forebrain organoids from each of the *NDE1* KO lines and the control line were split into two biological repeats (a total of 10 groups, 24 organoids in each group). The organoids were dissociated into single cells on day 32 using Accutase (A6964, Sigma), and the cell pellets were lysed with Tris-HCl NaCl buffer with 1% Triton X-100 to extract the proteins. Equivalent amounts of organoid cell lysates (50μg) were loaded onto a 10% SDS-PAGE gel. For experiments with Neural Stem Cell (NSC) cultures, 10 µg of protein lysate was loaded. The gels were subsequently electrophoretically transferred onto a Nitrocellulose membrane. To minimize any nonspecific interactions of the antibodies, the membrane was incubated for 1 h in a blocking solution (5 % non-fat milk powder in PBS-T, 1% PBS, and 1% Tween-20) at RT. After a brief wash with PBS-T, the membrane was incubated with primary antibodies overnight at 4°C and later washed with PBS-T three times for 5 min at RT. Primary antibodies included the NDE1 antibody (AP 10233-1, Proteintech, rabbit, 1:1000) to detect NDE1 and NDEL1 levels, and DM1A (Tubulin; T9026, Sigma, mouse, 1:1000) was used as a loading control. The NSC experiments included the following antibodies: mouse anti-pERK (Sigma M8159), rabbit anti-ERK (Sigma M5670), rabbit anti-Nde1 (Proteintech 10233-1-AP), mouse anti-LIS1^8^ (clone #338), rabbit anti-Lamin B1 (Abcam ab16048), and mouse anti-α-Tubulin (Sigma T9026, clone DM1A).

Subsequently, the membranes were incubated for 1h at RT with the secondary antibody at a 1:5000 dilution in the blocking solution (2.5% non-fat milk powder in PBS-T). Thereafter, the membranes were washed as described above. Antibodies bound to the target protein were detected using the ECL solution (20 ml HCL 8.5 pH, 44 µl p-coumaric acid, and 100 µl luminol). The secondary antibodies used are Peroxidase AffiniPure Goat Anti-Mouse or Rabbit IgG (H+L) from Jackson (115-035-003 or 111-035-144, respectively). Band intensities were quantified using Scuigo and ImageJ software. Statistical analyses were performed using GraphPad Prism. All the data were assessed for normal distribution and equal variance. For all hypothesis tests, exact test statistics, degrees of freedom, and confidence intervals were calculated are reported in the source data file. Exact P values were reported in the legends. All statistical tests were two-sided unless otherwise specified.

### On-chip brain organoids

On-chip brain organoids were cultured as previously described^3,9^. Briefly, the devices were fabricated using a commercial 6cm polystyrene tissue culture dish (Nunc). Eleven holes of 1.5mm in diameter were drilled through the dish bottom. A semi-permeable polycarbonate membrane (Whatman^®^ Nuclepore Track-Etched Membranes) was glued on the holes using a UV-curable adhesive (NOA81, Norland Products). The membrane covered nine of the holes, leaving two uncovered. The uncovered holes serve as inlets. A circular Polydimethylsiloxane (PDMS) stamp of thickness and diameter was placed on a 24x24mm^2^ microscope coverslip. The UV-curable adhesive was flown between the coverslip and the PDMS, thus forming a spacer with a thickness of 150μm. The spacer was half-cured by UV exposure, and the PDMS was peeled off the coverslip.

Approximately 900 cells were seeded per well into low-adhesion V-bottom 96-well plates to promote the formation of aggregates. Seventy-two hours post-seeding, the cell aggregates were transferred onto the fabricated culture dish. Nine aggregates were placed in each device on top of the membrane-covered holes. The device was sealed by adding the fabricated spacer-coverslip.

The plates were then filled with neural induction media, replaced every other day. Four days later, a collagen-laminin-based hydrogel (100% Matrigel) was injected into the device. Neural differentiation media supplemented with EGF (20ng/ml) and FGF2 (20ng/ml) were added to the device. Media was exchanged every other day.

For mitosis analysis (Fig. 1K-L), on-chip organoids were live-imaged every 2 minutes for several hours, eight days after Matrigel injection, with a 40x lens. Z-stacks (31 slices over 150 microns were acquired on a spinning disk confocal microscope (Andor Technology).

### MARS-Seq

Total RNA was extracted using the RNeasy Mini kit (Qiagen) under the manufacturer’s protocols. RNA was extracted from ESCs and on-chip organoids 18 days after embedding in Matrigel. Three repeats were taken. Organoid RNA was extracted directly from the device, with a total of 30 to 40 organoids per repeat and three repeats per experiment. RNA concentration and integrity were measured using Nanodrop (Thermo Scientific) and an Agilent Tapestation.

Libraries were prepared from up to 50 ng of total RNA. MARS-seq libraries were prepared as previously described^10^. The RNA was reverse-transcribed using barcoded oligo-dT primers with sample-specific index sequences and unique molecular identifiers. After first-strand cDNA synthesis, samples with similar cycle thresholds (CT) from quantitative PCR (qPCR) assessment were pooled, and the cDNA was converted into double-stranded DNA. This was followed by linear amplification through in vitro transcription using T7 RNA polymerase. The amplified antisense RNA was enzymatically fragmented and treated with DNase. The sequencing-ready libraries were created by ligating Illumina adapter sequences during the final reverse transcription step, followed by PCR enrichment and quality assessment using qPCR and the Agilent TapeStation system. The libraries were sequenced on an Illumina NextSeq 550 platform to obtain 75 bp single-end reads.

### Analysis of *NDE1* Expression Across the Cell Cycle

To investigate the dynamics of *NDE1* expression during the cell cycle, we analyzed single-cell transcriptomic data from the cortical excitatory neuron lineage. The UMAP embeddings, gene expression matrix, and cell type annotations were obtained from Braun et al^11^. Visualization of *NDE1* and *AURKA* gene expression across the UMAP embedding was performed using the scattern function from the *cytograph-shoji* package (https://github.com/linnarsson-lab/cytograph-shoji.git). To quantify expression patterns across distinct cell cycle phases, only cells from v3 Chromium chemistry samples were used. These cells were grouped according to their annotated cell cycle stage. Gene expression levels were summarized and visualized using the boxplot function from the *seaborn* Python library.

### RNA-Seq analysis

For MARS-seq, samples were analyzed using the UTAP pipeline^12^. Reads were trimmed and aligned to the GRCh38/hg38 reference genome for human using STAR (version v2.4.2a), with parameters –alignEndsType EndToEnd, outFilterMismatchNoverLmax 0.05, –twopassMode Basic. Gene read count was performed using the qCount function from the QuasR Bioconductor package (v.1.34)^13^ with default parameters. Gene quantification of the most 3’ 1000bp of each gene was performed using HTSeq-count^14^ in union mode while marking UMI duplicates (in-house script and HTSeq-count). Differential expression testing was performed with DESeq2^15^ (v.1.34) and pairwise comparison was performed with lfcShrink function with -type ashr^16^.

Genes with log2foldchange ≥ 1 and ≤ -1 with padj ≤ 0.05 and baseMean ≥ 10 were considered differentially expressed. Clustering was performed with the kmeans function in R.

### Real time-PCR

Total RNA was extracted using the RNeasy Mini kit (Qiagen) under the manufacturer’s protocols. RNA was converted to cDNA by qScript cDNA Synthesis Kit (Quantabio). Real-time PCR with SYBR FAST ABI qPCR kit (Kapa Biosystems) was performed using QuantStudio 5 Real-Time PCR System (Applied Biosystems). Each group contained three biological repeats, each with three technical repeats. If one of the technical repeats differs from the other two in more than one cycle, it is removed from the calculation. Thus, in each statistical group, n<=9. The delta CT values were used for statistical analysis, with *RPS29* as an internal control. Similar results were obtained with *GAPDH* as an internal control. The fold change was calculated using the delta-delta CT method, in which each delta CT value was normalized to the average CT value of the control line at the specific time point. Fold change=2^-1ΔΔCT^. Primers were taken from the primerBank database^17,18^ and are detailed in Supplementary Table 1.

### Forebrain organoids

Forebrain organoids were grown as previously described^19^. Briefly, about 9000 hPSCs per well were dispensed into low-adhesion V-shaped 96-well plates, and aggregates were formed.

Forebrain differentiation media, containing WNT inhibitor and TGFβ inhibitor, were exchanged every other day. On day 18, organoids were transferred to a 10-cm non-cell adhesive plate and cultured in a second forebrain media in suspension. Organoids were collected after two weeks, at day 32. Images were taken every other day.

### Cerebellar organoids

Cerebellar organoids were grown as previously described^20^. To initiate aggregate formation, ∼6,000 hPSCs were seeded per well into low-adhesion V-bottom 96-well plates in cerebellar differentiation medium containing a TGF-β inhibitor. On day 2, the media were supplemented with FGF2 (50 ng/ml). Five days later, the growth factors were omitted. Aggregates were treated on day 14, and FGF19 (300 ng/ml) was added. On day 21, organoids were transferred to a 10-cm non-cell adhesive plate and cultured in suspension in a second cerebellum media. Organoids were harvested on day 28. Images were taken every other day.

### Immunostaining

Control and *NDE1* KO forebrain organoids were immersed for 1 h in 4% PFA on days 18 and 32 DIV. After three washes with PBS, the organoids were cryoprotected overnight in 20% sucrose at 4°C. Organoids were embedded in Optimal Cutting Temperature (OCT) compound, sectioned to 12 μm thick slices, and stained with the following antibodies: phospho-Histone H3 (pH3; rabbit, 1:200, Sigma 06-570), Ki67 (mouse, 1:400, BD Biosciences 550609), cleaved caspase-3 (rabbit, 1:200, Cell Signaling 9661), and NeuN (mouse, 1:200, EMD Millipore MAB377).

Hoechst 33342 staining (Invitrogen H3570) was used to label the nuclei in 8.1 μM (or 5 μg/ml), final concentration from the stock solution in DMSO.

For EdU detection, organoids were incubated with 15 μM EdU for 1 hour at 37 °C prior to fixation. Detection was performed using the Click-iT™ EdU Flow Cytometry Assay Kit (Thermo Fisher Scientific, C10634). Image analysis was performed using Imaris software Imaris software (version 10.1.1, Bitplane, Zurich, Switzerland). Hoechst-positive nuclei were counted, and the percentages of NeuN-, Ki67-, EdU-, and pH3-positive nuclei were quantified relative to the total number of Hoechst-stained nuclei. For cleaved caspase-3, the expression area was measured using the color threshold tool in ImageJ software (NIH, Bethesda, MD, USA). The total area of each slice was defined by Hoechst staining, and the relative caspase-3–positive area was calculated accordingly.

### Neural Stem Cell (NSC) Culture and Nuclear/Cytosolic Fractionation

Wild-type and *NDE1* knockout hPSC lines (*NDE1 KO* #3, *NDE1 KO* #4) were seeded at 100,000–120,000 cells/well in 12-well plates pre-coated with Matrigel. Y-27632 ROCK inhibitor (10 µM) was added at plating and removed the following day. Cells were cultured in mTeSR+ medium (STEMCELL Technologies) for 6–8 days until cultures reached confluency. The medium was then replaced with basal medium (50% DMEM/F-12, 50% Neurobasal, GlutaMAX, NEAA, B-27 minus vitamin A, N2 supplements, penicillin/streptomycin, and 100 µM β- mercaptoethanol). The next day, cells were treated with hEGF (PeproTech, 10 ng/mL), FGF2- G3^21^ (prepared by the core protein unit at the Weizmann Institute, 100 ng/mL), or Phorbol 12- myristate 13-acetate (PMA, Sigma P8139, 100 nM) for the indicated durations. Cells were harvested on ice, washed once in cold PBS, and lysed in IP buffer (50 mM Tris-HCl pH 7.5, 100 mM NaCl, 1% Triton X-100, 4 mM MgCl₂, 0.1 mM DTT) supplemented with protease inhibitor cocktail (APExBIO K1007) and phosphatase inhibitors (10 mM sodium orthovanadate, 1 mM NaF, 10 mM β-glycerophosphate). Lysates were incubated on ice for 30 minutes with intermittent mixing and centrifuged at 14,000 × g at 4°C. Supernatants were collected and protein concentrations measured using a BCA assay kit (Thermo Fisher). Samples were mixed with 1/3 volume of 4× Laemmli buffer, boiled for 5 minutes, and stored at 80°C.

For nuclear/cytosolic fractionation of PMA-treated cells, cells were washed in cold PBS, scraped into Nuclear Isolation Medium (10 mM Tris-HCl pH 8.0, 250 mM sucrose, 25 mM KCl, 5 mM MgCl₂, 0.1 mM DTT, 0.1% NP-40, plus protease and phosphatase inhibitors), homogenized with a Dounce homogenizer, and centrifuged at 900 × g for 10 minutes at 4°C. The supernatant was collected as the cytosolic fraction. Pellets were resuspended in cold IP buffer and pulse vortexed vigorously for 30 minutes, then centrifuged at 17,000 × g for 10 minutes at 4°C. The resulting supernatant was collected as the nuclear fraction.

### NMC generation and culture Device fabrication

The microfluidic device was fabricated as previously described^22^. Briefly, the microfluidic device consists of a polydimethylsiloxane (PDMS) structure attached to a coverslip. The PDMS structure has three parallel microfluidic channels separated by micro-posts. The central channel will be filled with Matrigel with neural tube-like structures embedded in it. The PDMS structure was fabricated by first mixing PDMS curing agent and base polymer (Sylgard 184; Dow Corning) at a ratio of 1:10. The PDMS prepolymer mixture was cast onto a microfabricated silicon mold and baked at 110 °C for 1 h. Baked PDMS structure was punched with Harris Uni-Core punch tool (6 mm diameter, Ted Pella) to make medium reservoirs. PDMS stamps for microcontact printing were fabricated by casting PDMS prepolymer mixture onto a microfabricated silicon mold. The ratio between curing agent and base polymer was 1:20 for the stamps. This was baked at 110 °C for 1 h and 30 min. After baking, PDMS stamps were peeled off, sterilized using 100% ethanol, and coated with 1% (v/v) Matrigel solution at 4 °C overnight. The next day, glass coverslips (Thermo Fisher Scientific) were cleaned prior to ultraviolet ozone treatment. Coverslips were first sonicated in 2% (v/v) Hellmanex III (Hellma USA, 9-307-011-4- 507) for 30 min. Then, they were rinsed 3 times in deionized water, sonicated in 100% ethanol for 30 min, and blow-dried. Cleaned coverslips were treated with ultraviolet ozone (Ozone cleaner; Jelight) for 7 min. The Matrigel-coated PDMS stamps were blow-dried and stamped onto the ozone-treated coverslips to transfer Matrigel adhesive patterns onto the coverslips.

Stamps were peeled off and PDMS structures were attached to the coverslips.

### NMC culture

The NMC were generated similarly as previously described^22^. On day 0, hPSCs were dissociated from tissue culture plates using Accutase (Sigma-Aldrich) and resuspended at a concentration of 20 × 10^6^ cells per mL in mTeSR containing the ROCK inhibitor Y27632 (10 μM, Tocris) to prevent dissociation-induced apoptosis of hPSCs^23^. 10 μL of the hPSC suspension was injected into the central microfluidic channel. The microfluidic device was incubated for 45 min to allow hPSCs to attach to the Matrigel adhesive islands. Unattached cells were flushed out by adding fresh mTeSR containing 10 μM Y27632 to the central microfluidic channel. The next day, mTeSR medium was aspirated from the microfluidic channel, and 10 µL of 100% Matrigel was injected. The device was incubated for 10 minutes for gelation and establishment of a 3D culture environment. Right after gelation, neural induction medium (NIM) was added to both central reservoirs. NIM consisted of basal medium supplemented with TGF-β inhibitor SB431542 (10 μM, StemCell Technologies) and BMP inhibitor LDN193189 (1 µM, StemCell Technologies). The basal medium consisted of a 1:1 mixture of DMEM/F12 (Gibco) and neurobasal medium (Gibco) supplemented with 1% N2 supplement (Gibco), 2% B-27 supplement (with vitamin A, Gibco), Glutamax (2 mM, Gibco), 1% non-essential amino acids (Gibco), 100 µM 2-mercaptoethanol (Gibco), and 1% antibiotic–antimycotic (Gibco). From day 2 to day 4, CHIR99021 (CHIR, 3 μM, StemCell Technologies) and RA (250 nM, StemCell Technologies) were supplemented to the caudal reservoir, establishing rostral-caudal patterning. The rostral reservoirs of some samples were supplemented with either XAV939 (XAV, 5 µM, Cayman Chemical) or Phorbol 12-myristate 13-acetate (PMA, 200 nM, Selleckchem) or both.

From day 4 to day 9, all reservoirs were filled with NIM without supplementation, and the medium was changed every other day.

### ScRNA-Seq of forebrain organoids and NMC

Forebrain organoids from two control batches and *NDE1* KO lines 3 and 4 were collected at day 32 after aggregation and dissociated to single cells using Accutase (A6964, Sigma). Then, libraries were made according to Barcode technology for Cell Multiplexing protocol (CG000391 Rev B). The quality of the libraries was assessed by TapeStation and qPCR, and high-quality libraries were sequenced by the Nancy and Stephen Grand Israel National Center for Personalized Medicine (G-INCPM) using a NovaSeq 6000 sequencer.

To access the NMC, PDMS structures were manually detached from the coverslips. One device was used for the scRNA-Seq. The NMC was cut in half, and the rostral and caudal halves were collected separately. The collected tissues were dissociated into single cells by incubating in Accutase for 2 hr. The cell suspensions were centrifuged and resuspended in PBS containing 0.5% BSA and filtered through a 40 μm cell strainer (pluriSelect USA, 43-50040-51) to remove debris and aggregates. Dissociated and filtered cells were loaded into a 10X Genomics Chromium system (10X Genomics) within 1 hr. 10X Genomics v.3 libraries were prepared following the manufacturer’s instructions. Then, libraries were sequenced using paired-end sequencing with a minimum coverage of 20,000 raw reads per cell using an Illumina NovaSeq 6000. The scRNA-seq was performed as a service by the University of Michigan Advanced Genomics Core unit.

### scRNA-Seq analysis

Cell Ranger 7.0 was used to align the fastq read to the mm10 genome, filter and to generate gene-level unique molecular identifier counts (UMIs) and gene expression matrix. The expression matrix was then loaded into the R package Seurat (v4.0.0^24^), and cells suspected of being doublets or of low quality were removed. Specifically, for the cortical scRNAseq, we removed cells with >10% mitochondrial reads, cells with <100 and >10000 features, and cells with <200 and >30000 UMIs. For the rostral NMC scRNAseq, we removed cells with >10% mitochondrial reads, cells with < 200 and > 50000 features, and cells with < 1000 and >10000 UMIs. After quality control, the matrix was normalized and scaled with the sctransform algorithm. After PCA reduction, additional UMAP dimensionality reduction, clustering, and DEGs analysis. The Louvain algorithm was applied to cluster the cells, with 25 PCs selected and a resolution of 0.6 for the cortical organoids, and 30 PCs and 0.4 for the NMC tissues. The FindAllMarkers function in Seurat was used to calculate DEGs among different clusters, using the Wilcoxon rank sum test, minimum fraction of cells (in either group) > 0.4, and log2FoldChange > 0.5.

Cell Cycle Score was calculated using the function CellCycleScoring and the canonical gene sets for the S phase and G2/M phase. For the rostal organoids, the cell cycle score was used as a regression factor during the scaling. As this was not sufficient to remove the strong cell-cycle effect from the clustering, we removed 147 cell-cycle-related genes from the matrix, which appeared to be part of the top-loading genes in the first 5 PCAs. After the removal of those genes, the matrix was reanalyzed as explained above.

### Annotation of cortical organoid clusters

Brain region clusters (forebrain, midbrain/hindbrain) were assigned using marker genes described in Braun et al.^11^ (Supplementary Table 2). Oligodendrocyte precursor cells (OPCs), choroid plexus (CP), glia (Gl), ependymal cells (EP), neural crest cells (NC), Cajal Retzius cells (CR) (added marker genes from Pellegrini et al.^25^, layer VI (LVI) were annotated using PanglaoDB Augmented 2021, Azimuth Cell Types 2021, using Enrichr^26–29^.

### Mapping scRNA-seq cells to dissected regions in Braun et al.^11^

To assign regional identities to cells in cortical organoid samples, radial glia were extracted from the single-cell transcriptomic reference atlas provided by Braun et al.^11^ Cells annotated as "Radial Glia" were selected from all brain regions except those labeled "head" and "brain," which lacked defined regional information.

Radial glia from the 5 pcw sample were excluded, as many cells annotated as "forebrain" in this sample mapped to non-forebrain regions, suggesting a potential dissection error.

Organoid cells were mapped to the reference radial glia using Symphony^30^. Changes in regional cell-type abundance between conditions were assessed using a two-sided Fisher’s exact test.

Projection of NMC organoid transcriptome to human brain embryonic atlas^11^

To deduce the identity of NMC organoids, the transcriptome dataset was projected to human brain embryonic atlas. First, radial glia cells (5 and 5.5 PCW) were isolated from the atlas. Then, the NMC transcriptome was projected to the isolated human reference using

FindTransferAnchors^31^ and TransferData^31^ functions. MapQuery^31^ was used to project the NMC cells onto the human reference UMAP in Fig. 2K-O.

### Whole-Mount Immunohistochemistry

Pregnant females were assigned to a timed pregnancy schedule. At embryonic day 10.5 (E10.5), females were injected intraperitoneally with EdU at a dose of 0.05 mg/g body weight (prepared from a 30 mM stock in 1× PBS), 30 minutes prior to embryo dissection. Animals were euthanized via cervical dislocation, and embryos were dissected into cold PBS, washed once on ice in cold PBS, and fixed in 4% paraformaldehyde (PFA) in PBS for 20 minutes. Post-fixation, embryos were washed twice (5 min each) in cold PBS and subsequently immersed for 10 minutes each in 50% methanol/PBS, 70% methanol/PBS, and 100% methanol. Embryos were stored at −20°C until further processing. Staining was performed according to Yokomizo et al^32^. On the day of staining, embryos were rehydrated by sequential immersion in 75%, 50%, and 25% methanol/PBS for 10 minutes each. After two washes in PBS-T (0.4% Triton X-100), selected embryos underwent click chemistry for EdU detection using Cy5-Azide. Embryos were then blocked in 10% horse serum/0.1% BSA in PBS-T (0.3% Triton X-100) and incubated in click reaction buffer containing 0.1 M Tris-HCl (pH 8.5), 2.5 µM Cy5-Azide, 1 mM CuSO₄, and 0.1 M ascorbic acid. Following washes, embryos were blocked again in PBS-MT (1% w/v skim milk, 0.4% Triton X-100, 1× PBS) for 1–2 hours prior to overnight incubation with primary antibodies diluted in ½ PBS-MT (1.5% skim milk, 0.4% Triton X-100, 1× PBS) at 4°C on a shaker. For anti-OCT2 antibodies, incubation was extended to two overnights. Primary antibodies were washed twice in PBS-MT and once in PBS-T (1 hour per wash). Secondary antibody solutions were prepared using Alexa Fluor 488 or 555-conjugated donkey anti-mouse or anti-rabbit antibodies (Abcam) at 1:1000 dilution, along with 10 µg/mL Hoechst 33342 (Invitrogen H3570). Embryos were incubated overnight at 4°C in the secondary antibody solution with shaking. Post-incubation, embryos were washed three times in PBS-T and once in PBS (1 hour per wash), followed by a brief 10-minute post-fixation step in 4% PFA at room temperature. Embryos were washed three times in PBS and stored in PBS at 4°C for up to 3 days until imaging. For imaging, embryos were mounted in glass-bottom dishes (MatTek Life Sciences) and cleared with RapiClear 1.49 (SUNJIN LAB) for 10–30 minutes. Imaging was performed using a Dragonfly confocal microscope (Oxford Instruments, Andor) equipped with a Zyla cooled CCD camera. A 20× multi-immersion objective with glycerol as the immersion medium was used. Z-stacks were acquired at 10 µm intervals across 400 µm (41 sections), typically using 5×5 stitching across 3–4 fluorescence channels. Image stitching was performed with Andor Fusion software, and image analysis was carried out using Imaris (Bitplane) and Fiji (ImageJ, NIH).

### Whole-Mount in situ (Hybridization chain reaction, HCR)

Embryos were dissected in cold PBS, washed, and sequentially immersed in 50% methanol/PBS- T (0.1% Tween 20) and then in 100% methanol. Samples were stored at −20°C in 100% methanol until further processing. Hybridization chain reaction (HCR) was performed according to the protocol provided by Molecular Instruments with minor modifications. E10.5 embryos were photobleached using H₂O₂ and NaOH, as described in Morabito et al.^33^, followed by washes, proteinase K treatment, and immediate postfixation in 4% PFA on ice for 20 minutes.

### Embryos were washed with PBS-T

To prevent ventricular collapse due to high osmolarity, probe hybridization buffer was diluted in 5× SSCT to final concentrations of 10%, 25%, 50%, and 75%, with gradual progression to 100% (30 minutes per step). A similar gradient was used for the introduction of the amplification buffer. Hoechst 33342 (Invitrogen H3570) was added during the final 30-minute 5× SSCT wash. After amplification and washes, embryos were cleared in RapiClear 1.49 and imaged in MatTek glass-bottom dishes using an Andor Dragonfly system equipped with an Xtreme CCD camera.

Probes we used were: Lmx1b #NM_010725.3 (B3) and Pax2 #NM_011037.4 (B2) and the Amplifiers B3-594 and B5-647. Another set of probes used were: Dlk1 #NM_001190703.1 (B2) and En1 #NM_010133.2 (B4) and the Amplifiers B2-488 and B4-647.

## Supplementary Data S1. (separate file)

**Differentially expressed (DE) across genotype throughout development.** DE expressed genes from on-chip brain organoids comparing WT and *NDE1* KO from Day 18 (sheet 1, DE_genotype_D18), Day 5 (sheet 2, DE_genotype_D5), human pluripotent stem cells (sheet 3, DE_genotype_hPSC), and all combined (DE_across_genotype_development).

**Supplementary Fig. S1.**
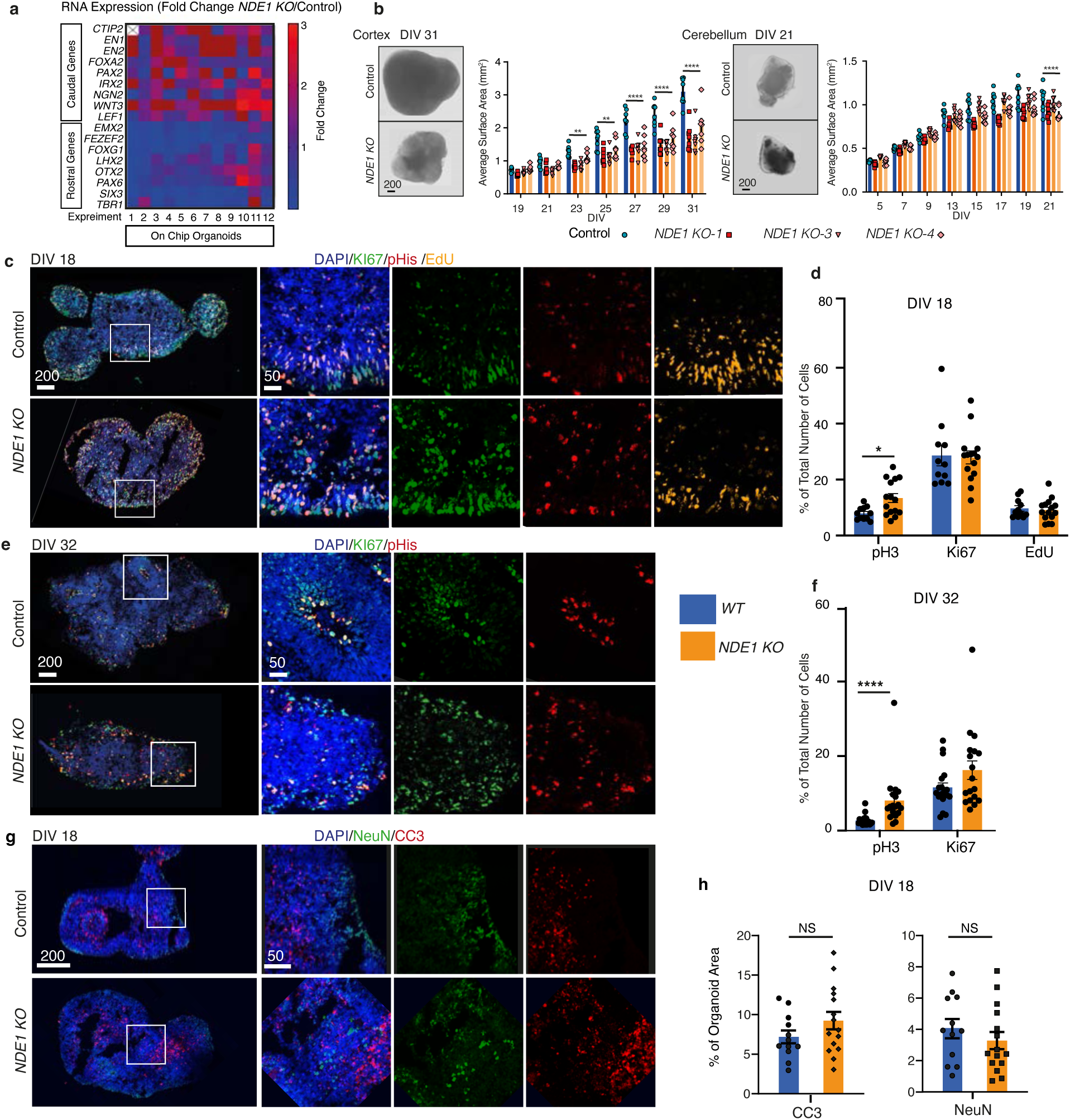
Additional characterization of proliferation and differentiation in *NDE1* KO brain organoids. (a) Heatmap neural differentiation gene expression marker (qPCR) comparing WT and *NDE1* KO organoids (on-chip organoids, 12 biological repeats). (b) Representative brightfield images and quantification of cortical and cerebellar organoid size (average surface area) in WT and *NDE1* KO organoids at indicated differentiation timepoints (days *in vitro*, DIV) mean ± SEM, n = 8 organoids per timepoint, *P* <0.0001). Scale bars: 200 µm. (c) Representative immunofluorescence images at early differentiation stage (DIV 18) showing proliferative (KI67, green), mitotically active (pHis, red), S-phase (EdU, yellow), and nuclear (DAPI, blue) markers in WT (control) and *NDE1* KO organoids. Scale bars: 200 µm (overview), 50 µm (insets). **(d)** Quantification of immunofluorescence markers on DIV 18 as percentages of total cells (mean ± SEM, n = 12-15 organoids per group; statistical test: **\*\*\*\***P < 0.0001*). **(e)** Representative immunofluorescence images at a later differentiation stage (DIV 32) showing the same markers in WT (control) and *NDE1* KO organoids as in **(c)**. Scale bars: 200 µm (overview), 50 µm (insets). **(f)** Quantification of cells positive for pHis or KI67 on DIV 32 as percentages of the total number of nuclei (mean ± SEM, n = 12-15 organoids per group; statistical test: **\*\*\*\***P < 0.0001*). **(g)** Representative immunofluorescence images at the early (DIV 18) differentiation stage, showing neuronal (NeuN, green), cell death (Cleaved Caspase 3, CC3, red), and nuclear (DAPI, blue) markers in WT (control) and *NDE1* KO organoids. Scale bars: 200 µm (overview), 50 µm (insets). **(h)** Quantification of respective immunofluorescence markers as percentages of total organoid area (mean ± SEM, n = 12-15 organoids per group; statistical test: **\*\*\*\***P < 0.0001*).

**Supplementary Fig. S2:**
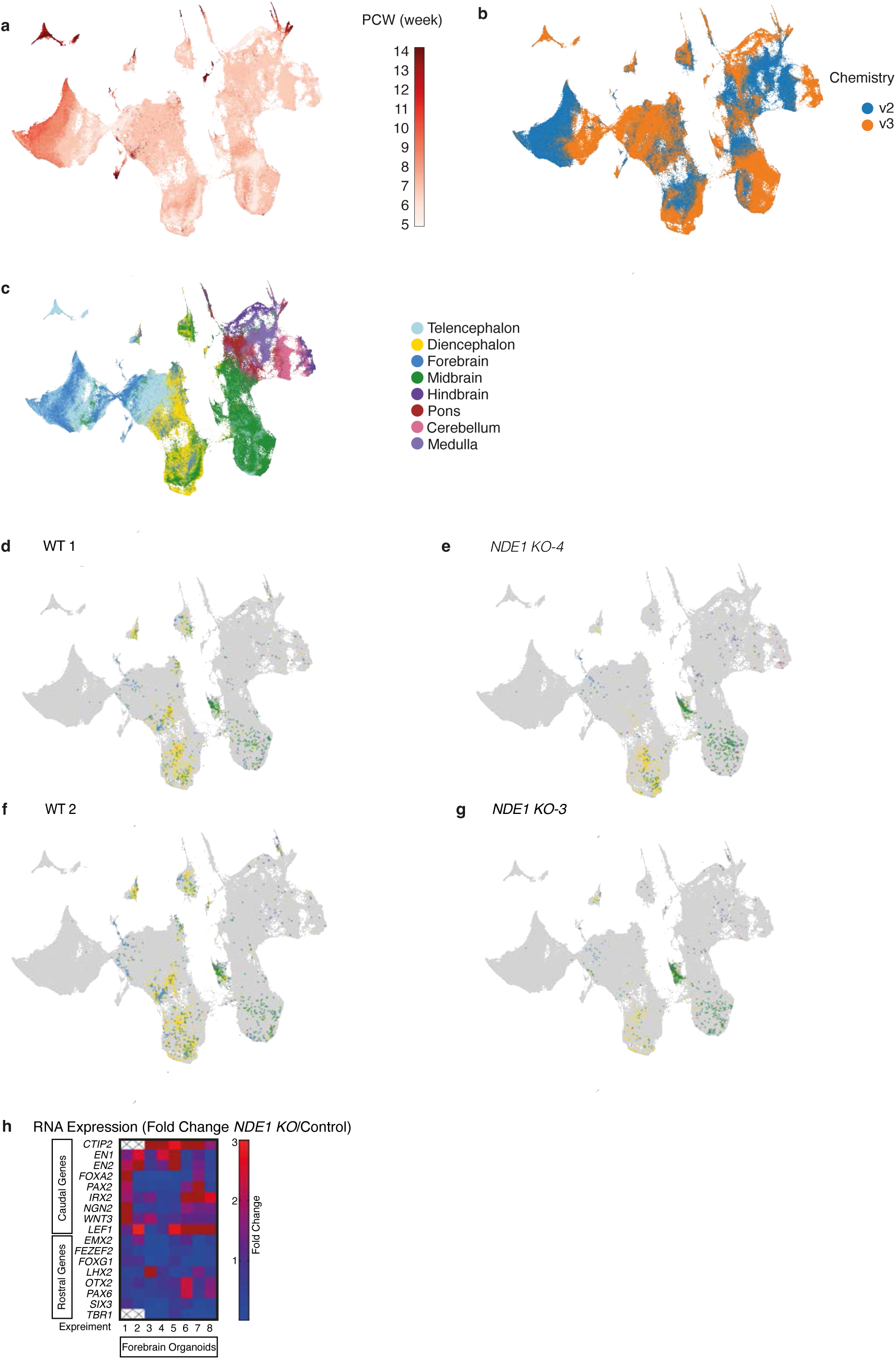
UMAPs of scRNA-Seq data of human cortical organoids. (a-b) UMAP visualization of radial glia cells from the human embryonic brain atlas^11^, colored by (**a**) developmental age in post-conceptional weeks (PCW) **(b)** Genomics Chromium chemistry version (v2 and v3) (**c**) UMAP visualization of radial glia cells from the human embryonic brain atlas^11^(PCW 5.5), color coded by anatomical region. **(d-g)** Cortical organoid cells mapped onto the radial glia cells from the human embryonic brain atlas^11^. Cells are colored by their assigned regional identity based on the mapping; reference atlas cells are shown in gray. **(d, f)** Wild-type (WT) organoid cells. **(e,g)** *NDE1* KO organoid cells. **(h)** Heatmap neural differentiation gene expression marker (qPCR) comparing WT and *NDE1* KO forebrain organoids, eight biological repeats.

**Supplementary Fig. S3:**
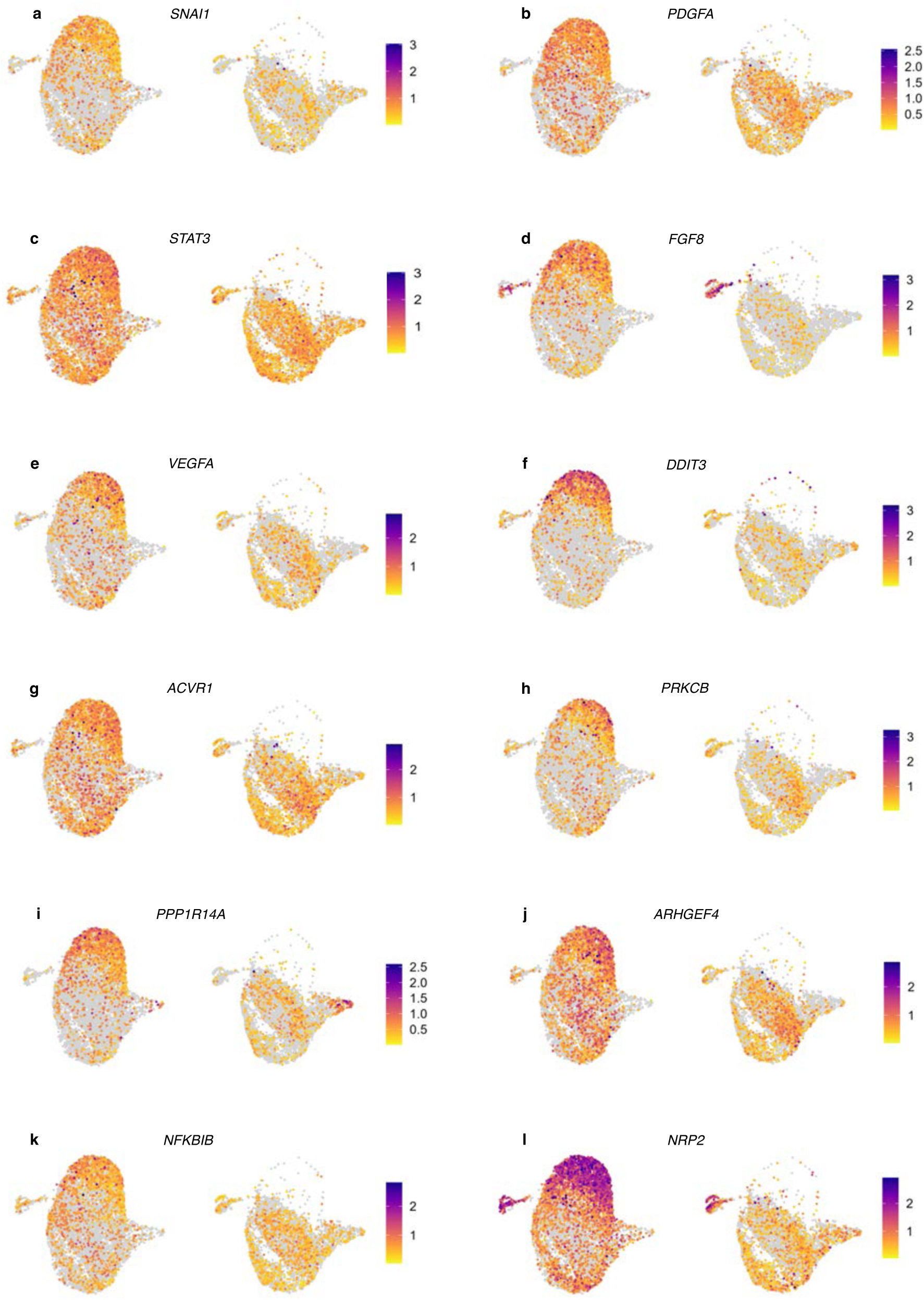
UMAP visualizations of representative genes linked to ERK signaling, showing differential expression between WT and *NDE1* KO NMC (expression scale indicates normalized expression levels per gene). *SNAI1* (**a**), *PDGFA* (**b**), *STAT3* (**c**), *FGF8* (**d**), *VEGFA* (**e**), *DDIT3* (**f**), *ACVR1* (**g**), *PRKCB* (**h**), *PPP1R14A* (**i**), *ARHGEF4* (**j**), *NFKBIB* (**k**), *NRP2* (**l**).

**Supplementary Fig. S4.**
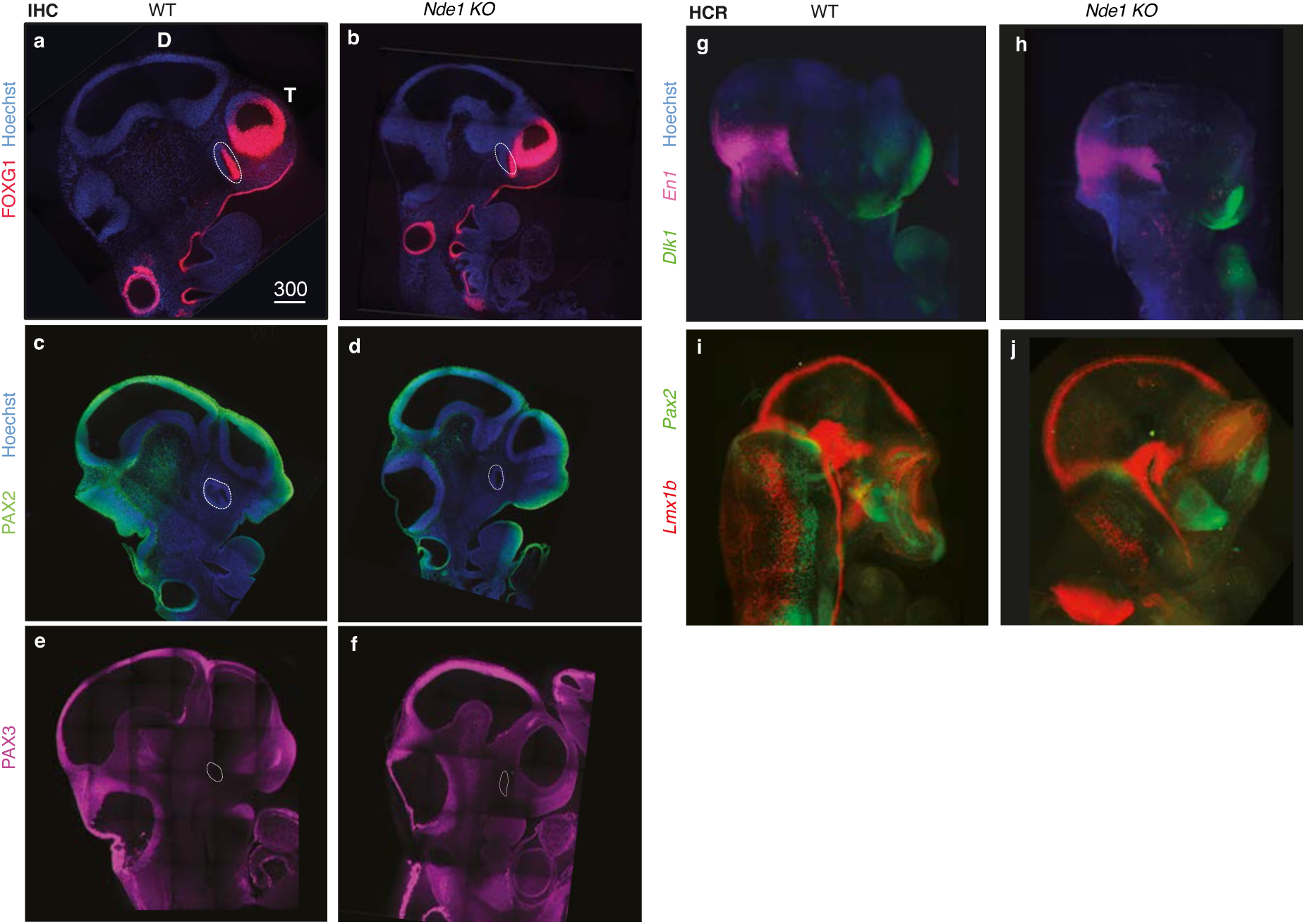
Expression patterns of multiple regional markers remain largely unaffected in *Nde1* KO embryos. (**a–f**) Immunohistochemical staining comparing FOXG1 (**a**, **b**), PAX2 (**c**, **d**), and PAX3 (**e**, **f**) protein expression between WT and *Nde1* KO E10.5 mouse embryos. Hoechst (blue) stains nuclei. d, diencephalon; t, telencephalon. Scale bar (**a–f**) 300 µm. (**g–j**) Hybridization Chain Reaction (HCR) for RNA transcripts of indicated markers: *En1* and *Dlk1* (**g**, **h**), and *Lmx1b* and *Pax2* (**i**, **j**) in WT and *Nde1* KO embryos. Hoechst (blue) nuclear staining is included in panels **g** and **h**. Scale bars: 300 µm. Representative images are from n=3 embryos per genotype.

**Table S1.**
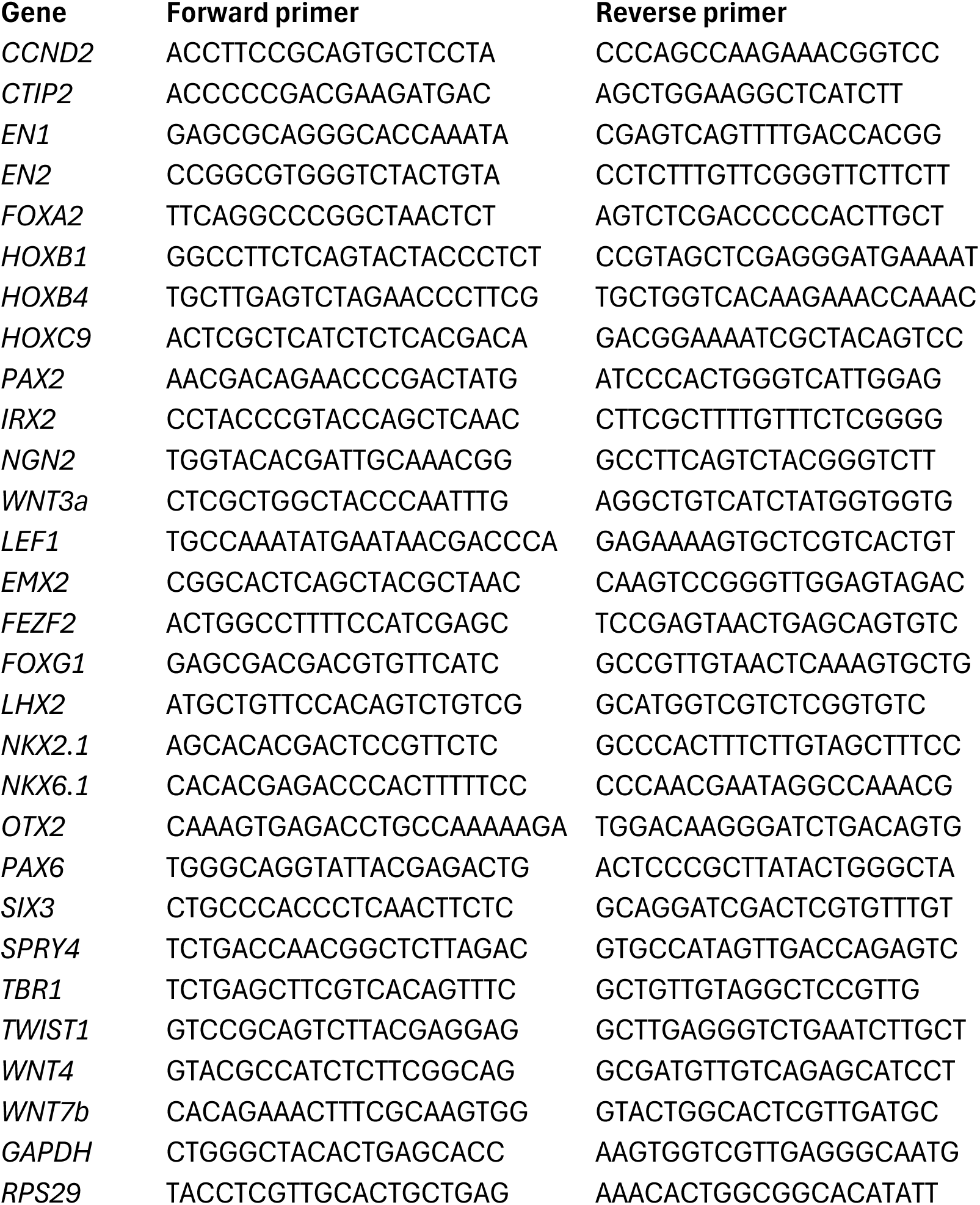
List of primers used for real-time PCR.

**Table S2.**
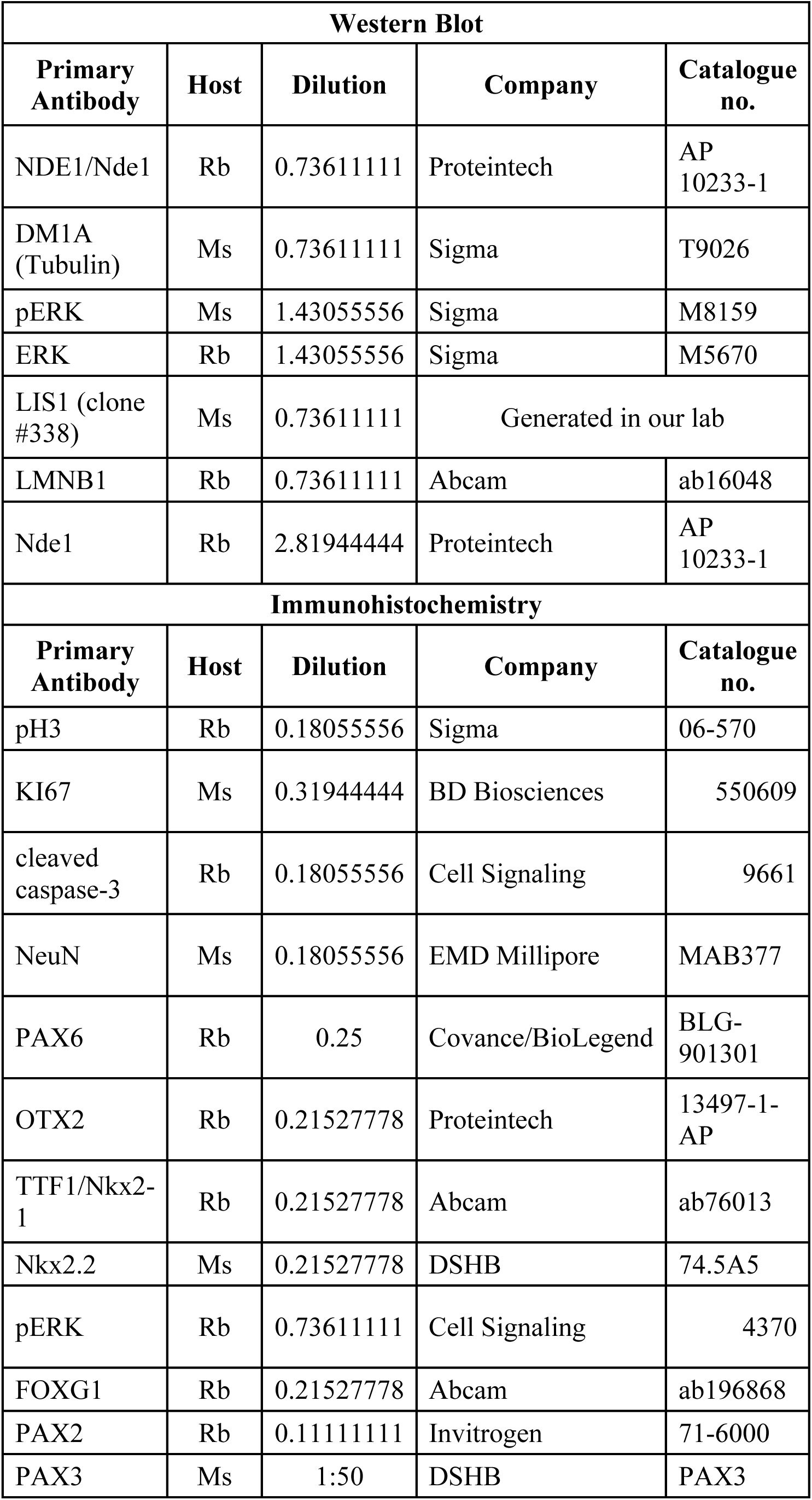
List of antibodies used in the study.

